# Historical texts as a potential resource for plant-derived natural products against SARS-CoV-2 – the example of the *Receptarium* of Burkhard III von Hallwyl from 16th century Switzerland

**DOI:** 10.1101/2024.06.13.598798

**Authors:** Nina Vahekeni, Jonas Stehlin, Corinna Urmann, Evelyn Wolfram, Yannick Geissmann, Yelena Ruedin, Olivier Engler, Andreas Lardos

## Abstract

In the search for more effective prophylactic and possibly curative therapeutics against SARS-CoV-2, an historical-ethnobotanical approach was used to select plants described in the *Receptarium* of Burkhard III von Hallwyl (RBH), an influential recipe text from 16th century Switzerland. Ten species were identified based on specific historical uses presumably linked with the treatment of viral infections as well as inflammatory conditions. For each plant candidate, aqueous and hydroethanolic extracts have been produced. CellTiter-Glo® Luminescent Cell Viability Assay was used to assess antiviral activity against SARS-CoV-2 and the effect on cell viability of the extracts.

Of the ten plant species tested, four displayed an antiviral activity ≥ 50% at 16.7 µg/ml with acceptable cell viability (> 75%): *Sambucus nigra* L. (leaves), *Viola odorata* L. (leaves), *Geranium robertianum* L. (arial parts) and *Artemisia vulgaris* L. (aerial parts). The crude extracts were partitioned in aqueous and organic fractions and further analyzed. The ethyl acetate fractions of *S. nigra, V. odorata* and *G. robertianum* expressed significant antiviral activity of nearly 100% at 5.6 µg/ml (P < 0.05). The most potent inhibitory activity was observed for the ethyl acetate fraction of *Viola odorata* L. (leaves) with 87% at 1.9 µg/ml (P < 0.0001). Alongside bioactivity analysis phytochemical fingerprints were made, with the aim to understand important substance classes contained. Further investigations are required to explore the active principles.

Our study shows that an ethnopharmacological approach based on historical records of traditional use to select potential herbal candidates coupled with a rational screening process enables an efficient search for plant-derived natural products with antiviral activity against SARS-CoV-2.

## 1 Introduction

The global health impact of the coronavirus disease 2019 (COVID-19) pandemic, caused by the infection with severe acute respiratory syndrome coronavirus 2 (SARS-CoV-2), has been significant. The infection has caused over 6.9 million deaths across the globe by late December 2023 (WHO Coronavirus (COVID-19) Dashboard, https://covid19.who.int/). In addition, 15% of acute symptoms persist in a long-COVID, characterized among others by fatigue, dyspnea and mental disorder lasting several weeks after an acute COVID-19 episode (Sudre et al., 2021). Currently there are few therapeutical options for SARS-CoV-2 infection, with nirmatrelvir/ritonavir (Paxlovid^TM^) remaining the recommended first-line curative therapy in Switzerland to prevent a severe course of the COVID-19 disease (Swiss Society of Infectious Diseases (SSI), 2023). Remdesivir (Veklury®) has been authorized for emergency use in certain countries. In term of prevention, vaccination serves as the primary preventive measure and few drugs have shown effective prophylaxis (Sharif et al., 2021), notably monoclonal antibody products. However, the emergence of new circulating variants undermines the efficacy of monoclonal antibodies, which are specific for a given variant (Stadler et al., 2023). Limited treatment possibilities have resulted in a persisting global demand for therapy options, such as plant based complementary remedies.

Potent anti-coronavirus activities (COVID-19, SARS-CoV, MERS-CoV) of natural products have been summarized in several reviews (Huang et al., 2020; Verma et al., 2020; Attah et al., 2021; Kim, 2021; Llivisaca-Contreras et al., 2021; Remali and Aizat, 2021; De Oliveira et al., 2022; Khuntia et al., 2022; Latha et al., 2022). Various phytochemicals including polyphenols, alkaloids and terpenoids were identified as potential active anti-SARS-CoV-2 agents (Adhikari et al., 2021). Compared with the large majority of investigations evaluating the *in silico* potential of plant metabolites, few studies reported *in vitro* anti-SARS-CoV-2 activity of plant extracts (Nair et al., 2022; Chen et al., 2023; De Araujo et al., 2023; Farooq et al., 2024; Kakimoto et al., 2024; Rodrigues et al., 2024). Certain plants with a long history of use in traditional medicine systems to manage flu syndromes, as for example *Withania somnifera* (L.) Dunal, *Andrographis paniculata* (Burm.F) Wall.ex Nees, *Echinacea purpurea* (L.) Moench, *Artemisia annua* and *Nigella sativa* L., have shown to be promising candidates in the search for an effective preventive or curative treatment of COVID-19 (Kolev et al., 2022; Nair et al., 2022; Nicolussi et al., 2022; Cyril et al., 2023; Ramli et al., 2023; Shanker et al., 2023).

An editorial by (Wang et al., 2021) in this journal commented on the research topic “ethnopharmacological responses to the corona virus disease”, illustrating the enormous potential of treatment options for this health problem from traditional medicine systems, but also pointing out the challenges faced by research in this context.

In search for effective agents for the potential development of prophylactic or therapeutic treatment options against SARS-CoV-2, we pursue the question of whether traditional knowledge from past eras that has been handed down in writing can offer starting points for research in this regard. In recent decades, the study of historical texts has attracted research interest in ethnopharmacology (Lardos, 2015). A prominent example illustrating such potential was the development of artemisinin from *Artemisia annua* L. (Asteraceae) as a medication for malaria, which was awarded with the Nobel prize in 2015. Its discovery goes back to the study of a 4th century CE text of Chinese medicine (Hsu, 2006).

Here we focus on the *Receptarium* of Burkhard III von Hallwyl (RBH), an early New High German recipe text from Switzerland dated to 1580 (Frei, 2012). The text belongs to the genre of “Hausarzneibücher”, whose target audience consisted primarily of medical laypeople (Fankhauser, 2012). This is also evident from the text’s preface, which mentions that the work is dedicated to the “common man”. The fact that up to the 18th century at least 15 manuscript copies were produced and distributed in various regions of Switzerland suggests that RBH must have had a remarkable local significance (Frei Haller, 2005). In a recent study by Stehlin et al. (2024), the dermatological recipes contained in the text were analyzed and cataloged in a relational database comprising 196 recipes which describe the use of 156 plants for 52 different uses for skin complaints, wounds or skin manifestations. We use the RBH database of dermatological recipes as a source for the selection of candidate plants for *in vitro* screening regarding their antiviral potential.

## 2 Materials and Methods

### 2.1 Selection of candidate species

Candidate species were selected from the RBH database of dermatological recipes in Stehlin et al. (2024) based on the following criteria. The respective RBH plant name, i) appears in plant use records in which at least one of the interpretations of the historical use leads to infectious diseases of viral origin, the therapy of which consequently requires medication with antiviral properties (among others), as specified in Table 1 in Stehlin et al. (2024); ii) appears in plant use records mentioned in recipes with only one active ingredient of plant origin (herbal simples). This allows to ascribe any potential bioactivity observed to one specific ingredient. For details of the methodology regarding the interpretation of the medicinal uses mentioned in RBH and the assessment of the RBH plant names, see Stehlin et al. (2024).

**Table 1:**
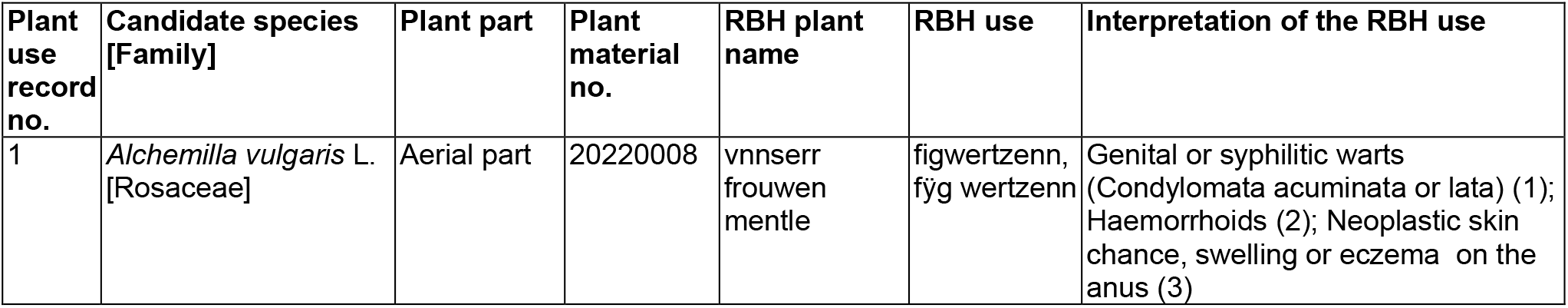
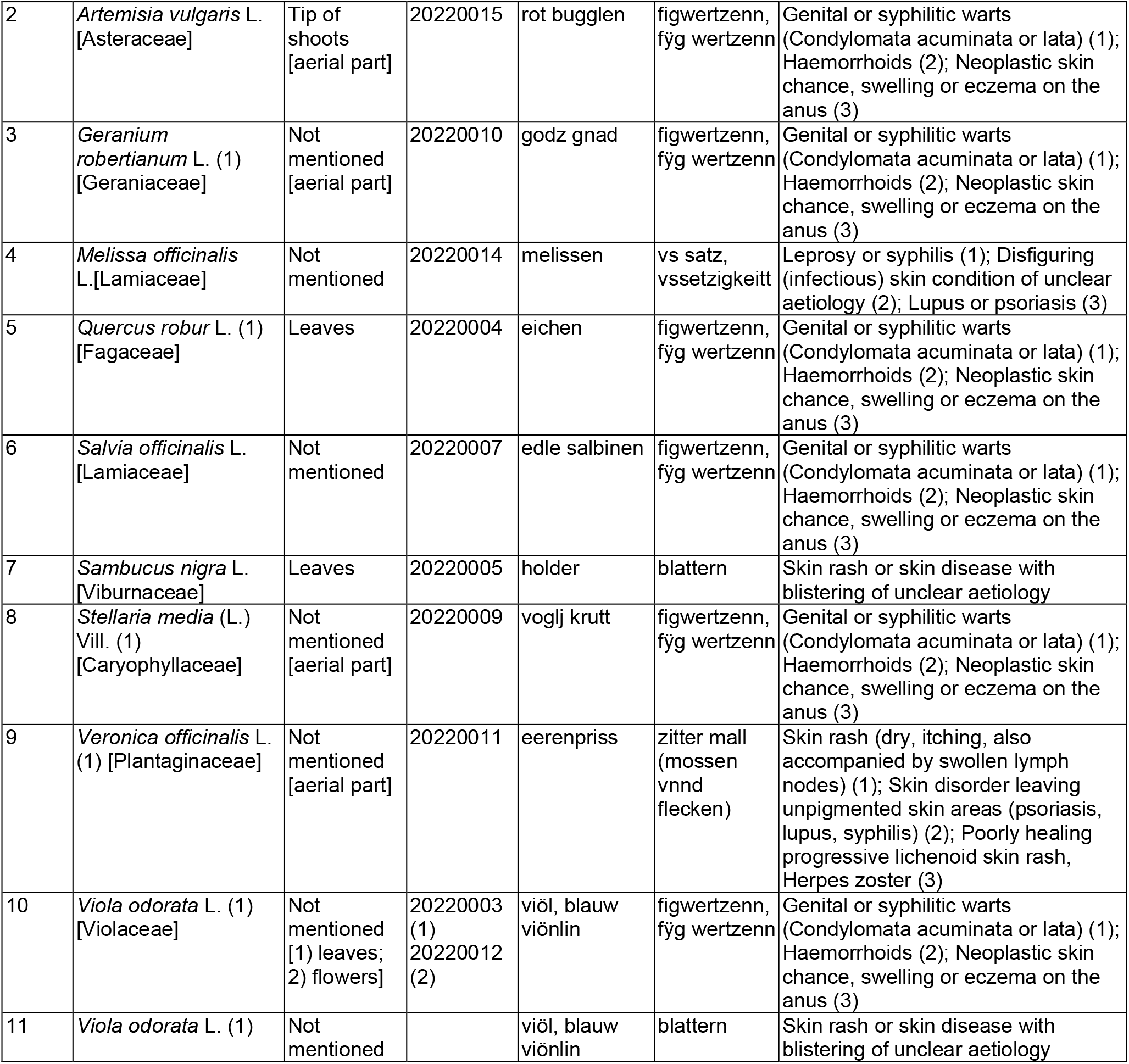
Candidate species selected from the RBH database of dermatological recipes in Stehlin et al. (2024). Candidate plant: Botanical identity of the RBH plant name (*). Only candidate species with the highest chance of being the correct attribution were considered (based on information in Stehlin et al. (2024, Table 1). Plant part: Plant part used as mentioned in the respective RBH recipe (*). Plant parts of purchased samples differing from the plant parts mentioned in the respective RBH recipes are indicated in square brackets. Plant material no.: ZHAW in-house material number of purchased samples of candidate plants. RBH plant name: Plant name mentioned in RBH (*).RBH use: Medicinal use mentioned in the respective recipe in RBH (*). *(Data was adopted from Stehlin et al. (2024, Tables 1 and 2).

### 2.2 Plant supply and extracts preparation

Dried plant material was purchased from Dixa AG in Switzerland (for detailed information, see **Supplementary Material, Table S1**). As far as possible the same plant parts of the candidate species as mentioned in the respective RBH plant use record was sourced from the supplier. Two different types of extraction procedures were carried out on the powdered dry material. The aqueous crude extract was prepared by adding a 20-fold quantity of water in relation to plant material and boiled for 15 min. The decoction was filtered with a Büchner funnel under vacuum or with a filter paper. The filtrates were freeze-dried and stored at -20 °C until use. For hydroalcoholic extraction, an 80% ethanol extract (EtOH 80%) was prepared by adding a ten-fold quantity of solvent in relation to plant material and extracted at room temperature for 2 h under constant agitation. Extracts were filtered through a folded filter paper, concentrated on a rotavapor (Büchi, Switzerland) at 40 °C until 60 mbar, freeze-dried, and stored at -20 °C until use. Preselected crude extracts were partitioned by liquid-liquid extraction using water and ethyl-acetate (1:1 v/v) in a separating funnel, allowing to recover two fractions, the aqueous fraction (AF) and the ethyl-acetate fraction (EAF). Detailed information on chemicals and materials is available in **Supplementary Material, Table S2**.

### 2.3 Screening procedure

Twenty-two crude extracts (11 aqueous and 11 hydroethanolic extracts) were tested for their anti-SARS-CoV-2 activity. Antiviral activity (inhibition of virus mediated cytopathic effect) and cell viability (absence of toxic effect) were assessed on Vero E6 cells. The crude extracts were screened in duplicate against SARS-CoV-2 using the CellTiter-Glo® Luminescent Cell Viability Assay. Crude extracts demonstrating an inhibitory activity ≥ 50% at a final concentration of 16.7 µg/mL with acceptable cell viability (>75%) were retained for a further fractionation step and underwent partition chromatography. The inhibitory activity and cell viability of the obtained fractions (AF and EAF) were assessed.

### 2.4 Cells and pathogenic virus

Vero E6/TMPRSS2 were passaged in Dulbecco’s Modified Eagle Medium (DMEM) containing 10% Foetal Calf Serum (FCS), 100 U /mL of penicillin, 100 μg /mL of streptomycin and 1mg/mL geneticin (all from Bioswisstec). SARS-CoV-2 (2019-nCoV/IDF0372/2020) was propagated in Vero E6/TMPRSS2 cells in DMEM containing 2% FCS, supplements (2%-FCS-DMEM) and 1 mg/mL geneticin at 37 °C, >85% humidity and 5% CO_2_. Viral titer was determined by standard plaque assay, by incubating ten-fold serial dilutions of the virus for 1 hour at 37 °C on a confluent 24-well plate with Vero E6/TMPRSS2 cells. Then, inoculum was removed, and 1 mL of overlay medium (20 mL DMEM, 5 mL FCS, 100 U /mL of penicillin, 100 μg /mL of streptomycin and 25 mL of Avicel rc581) was added. After 3 days of incubation at 37 °C, the overlay was removed, and the plates were stained with crystal violet. Due to the clearer read out, Vero E6 cells in suspension were used for the antiviral assays. Vero E6 cells were passaged in Minimal Essential Medium (MEM) containing 10% FCS and supplements (2 mM L-glutamine, non-essential amino acids, 100 U /mL of penicillin, 100 μg /mL of streptomycin and 1,5 g/L sodium bicarbonate, all from Bioswisstec) at 37 °C, > 85% humidity and 5% CO_2_.

### 2.5 Antiviral activity testing

Lyophilized plant extracts were resuspended in DMSO or H_2_O to 25 mg/mL and diluted to 400, 200, 66.6, 22.2 and 7.4 µg/mL in MEM containing 2% FCS and supplements (2 mM L-glutamine, non-essential amino acids, 100 U /mL of penicillin, 100 μg /mL of streptomycin and 1.5 g/l sodium bicarbonate, all from Bioswisstec) (2%-FCS-MEM). The plant extracts, 50 µL of each concentration, were distributed in duplicates in the upper half of a flat bottom 96 well plate (TPP, Trasadingen, Switzerland) to assess antiviral activity, and 50 µL of the same concentrations of plant extracts were distributed in duplicates in the lower half of the 96 well plate to assess cell viability or toxicity of extracts. On each plate 10 wells were used to determine maximum viral cytopathogenicity (virus control VC) of infected but untreated cells and 10 wells to determine maximum cell viability (cell control CC) of uninfected and untreated cells. Further controls included serial dilutions of remdesivir as positive control and DMSO at the same concentration as used for the extracts to control for effects of the diluent. Plates were transferred to the BSL-3 laboratory and 100 PFU SARS-CoV-2 virus (2019-nCoV/IDF0372/2020) in 50 µL culture medium (2%-FCS-DMEM) was added to the upper half of the plate as well as to the wells foreseen for the virus control VC, while to the lower half of the 96 well plate and to the cell control 50 µL culture medium (2%-FCS-MEM) was added. Plates were incubated for 1 h at 37°C and 5% CO_2_ and 100 µL of Vero E6 cell suspension (2 x 10e5 /mL) in cell culture medium (2%-FCS-MEM) was added to each well leading to a final concentration of extracts of 100, 50, 16.7, 5.6 and 1.9 µg/mL). Plates were then incubated for 72 h at 37°C and 5% CO_2_ and cell viability determined by CellTiter-Glo Luminescent Cell Viability Assay (Promega). In short, CellTiter Glo suspension was prepared according to the manufacturer’s protocol (and mixed 1:1 with cell culture medium (2%-FCS-MEM). The cell culture medium from incubated plates was removed and 200 µL of CellTiter Glo/medium mix added to each well. Plates were shaken for 2 min at 400 rpm incubated for 10 min at RT and 100 µL of the CellTiter Glo mix transferred to a 96 well half area white flat bottom plate (Corning ® Costar® 3693, Fisher Scientific). Luminescence was measured using the GloMax instrument (Promega).

### 2.6 HPLC-MS chemoprofiling

The ethyl acetate extracts were solved in methanol (analytical grad, VWR Chemicals, Darmstadt) in a concentration of 10 mg/mL and 5 µL of extracts were injected into a Shimadzu HPLC system consisting of two pumps (LC-20AD), an autosampler (SIL-20AC HT), a column oven (CTO-20A), and a PDA detector (SPD-M20). The HPLC method was based on a reversed-phase column (Phenomenex Kinetex C_18_ 2.1 × 100mm 2.6 u) and solvents A (water + 0.1% formic acid) and B (acetonitrile + 0.1% formic acid), with a flow of 0.4 mL/min and an oven temperature of 30°C. Separation was achieved using a gradient starting with a hold of 15% solvent B for 1 min, raising to 95% B within 35 min, and a hold of 95% B for 15 min. HRESIMS was recorded in positive ion mode (interface voltage 3.5 kV) and in negative ion mode (interface voltage = − 3.0 kV) using an IT TOF (Shimadzu). The interface was adjusted to T = 230°C and a nebulizer N_2_ stream of 1.5 L/min was used. The method included MS^1^ with m/z 100–1000 and ion accumulation of 10 ms as well as for MS^2^ with m/z 50–1000 and an ion accumulation 30 ms. CID energy was set to 50%. For data analysis LCMS Solution Version 3.80.410 (Shimadzu) and the implemented Formula Predictor Version 1.12 were used.

### 2.7 Data analysis

Percent cytopathic effect (CPE) inhibition and percent cell viability were calculated according to (Severson et al., 2007). Percent CPE was calculated from luminescence signal of [(plant extract – virus control) / (cell control – virus control)] * 100. Percent cell viability was defined as luminescence signal at (plant extract / cell control) * 100 at the extract final concentration.

The means of the inhibitory activity from all samples were compared to the relevant solvent control at each respective dosage using two-way ANOVA, with α = 0.05. Multiple comparisons were corrected using Dunnett’s method. All statistical testing was carried out using GraphPad Prism version 10.1.2 for Windows, GraphPad Software, Boston, Massachusetts USA.

## 3 Results

### 3.1 Plant candidates selected from RBH

Ten candidate species from 9 different botanical families involved in 11 plant use records were selected from RBH following the procedure described in 2.1 (**Table 1**). In each of the 11 use records at least one of the possible interpretations of the historical use in question points to an underlying viral disease. These include genital warts and herpes zoster but perhaps also specific cases of neoplastic skin changes or ill-defined but obviously infectious skin diseases that might be caused by a virus (**Table 1**). Based on the analysis in Stehlin et al. (2024, Table 1), any pharmacological intervention for the treatment of these conditions includes medication with antiviral properties. Consequently, the RBH plant mentioned in the record would be required to have antiviral properties in order to exert beneficial effects on the interpreted historical use. This background provided the rational basis for the selection of the ten plant species investigated in this study.

### 3.2 Anti-SARS-CoV-2 Screening

A total of 22 extracts were prepared from different parts of the 10 plant species using water and 80% ethanol as extraction mediums. A detailed description of plant species, plant parts extracted, solvents, drug-solvent ratio, and extraction yields is given in **Supplementary Material, Table S3**. All plant extracts were tested for their anti-SARS-CoV-2 activity. Antiviral activity and cell viability were assessed on Vero E6 cells in a serial diluted manner – 100, 50, 16.7, 5.6 and 1.9 µg/mL. Of the 22 crude extracts screened, only the hydroethanolic extracts showed an inhibitory activity. Four hydroethanolic active extracts, namely *Artemisia vulgaris* (ArV, aerial part), *Viola odorata* (ViO, leaves), *Geranium robertianum* (GeR, aerial part), *Sambucus nigra* (SaN, leaves), displaying an inhibitory activity ≥ 50% at a final concentration of 16.7 µg/mL were retained for further screening (**Figure 1, A**). At this concentration point, all four crude extracts showed a cell viability > 75% (**Figure 1, B**). The antiviral activity of the crude extracts at 16.7 µg/mL ranged from 65% for *A. vulgaris* to 97% inhibition for *S. nigra* (leaves).

**Figure 1:**
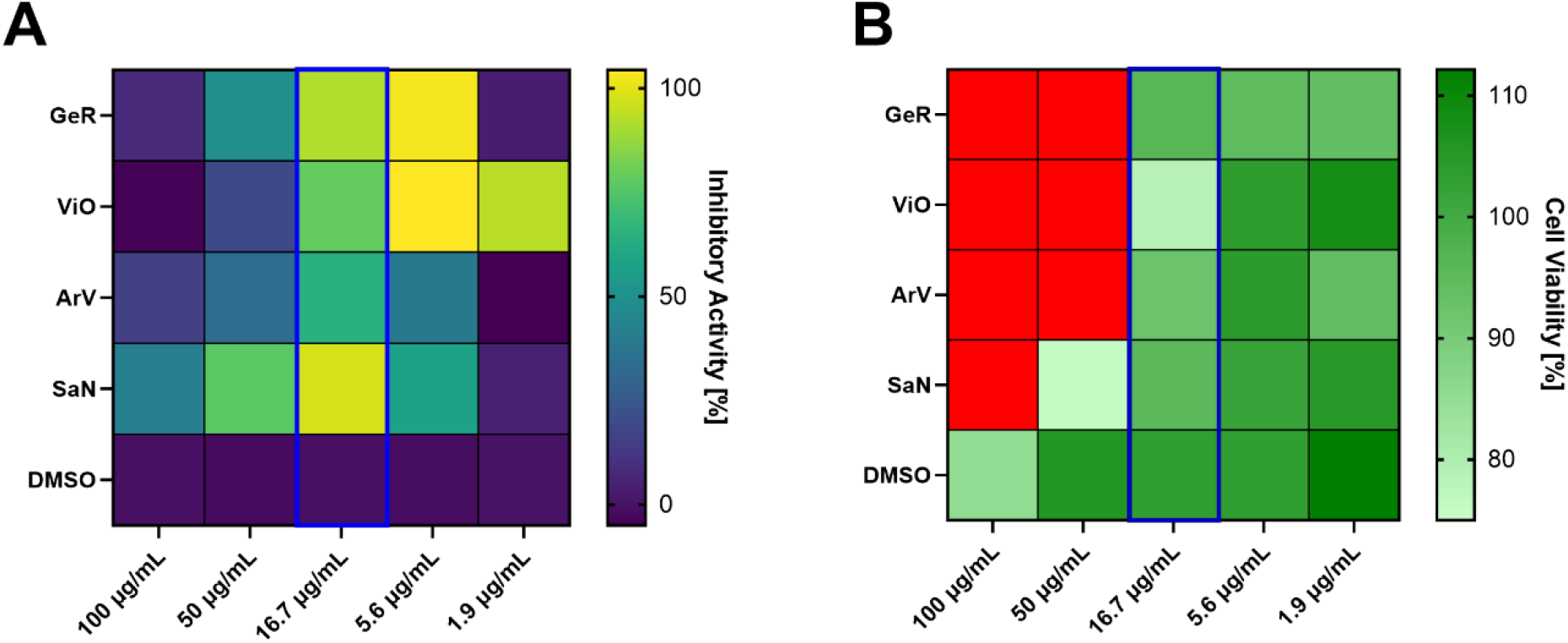
Crude extracts (A) inhibitory activity and (B) cell viability assessment. The four hydroethanolic crude extracts demonstrating an inhibitory activity ≥ 50% at a final concentration of 16.7 µg/mL and cell viability > 75% are represented: GeR - Geranium robertianum, ViO - Viola odorata, ArV - Artemisia vulgaris, SaN - Sambucus nigra A: Percent inhibition of cytopathic effect (CPE) is expressed as percent of inhibitory activity. Inhibitory activity data represent the mean of two independent replicates (n=2): ArV (65%), ViO (78%), GeR (92%), SaN (97%). Controls: DMSO and Remdesivir. Remdesivir was tested at a concentration of 10 µM and cell viability was 98%. B: Hydroethanolic crude extracts showing cell viability > 75% at final concentration of 16.7 µg/mL are represented: Green shades represent a cell viability ≥ 75%. Red color: value of cell viability < 75%.

### 3.3 Active organic fractions

The four hydroethanolic active extracts of *A. vulgaris* (aerial part), *V. odorata* (leaves), *G. robertianum* (aerial part) and *S. nigra* (leaves) were partitioned in organic and aqueous fractions via liquid-liquid extraction. Among aqueous fractions, *A. vulgaris* (ArV / AF) was the only one exhibiting significant inhibitory activity at 50 µg/mL (**Figure 2, A**). The ethyl acetate fractions (EAF) demonstrated promising anti SARS-CoV-2 activity at a final concentration of 5.6 µg/mL with a nearly 100% inhibitory effect for *A. vulgaris* (100%), *V. odorata* (100%) and *G. robertianum* (98%) (P < 0.05) (**Figure 2, B**). In parallel, these three organic fractions showed a reliable cell viability for the two lowest concentrations points(P < 0.05). The EAF of *V. odorata* exhibited the most potent antiviral activity, where 1.9 µg/mL reduced the virus mediated cytopathic effect by more than 85% (**Figure 2, B**).

**Figure 2:**
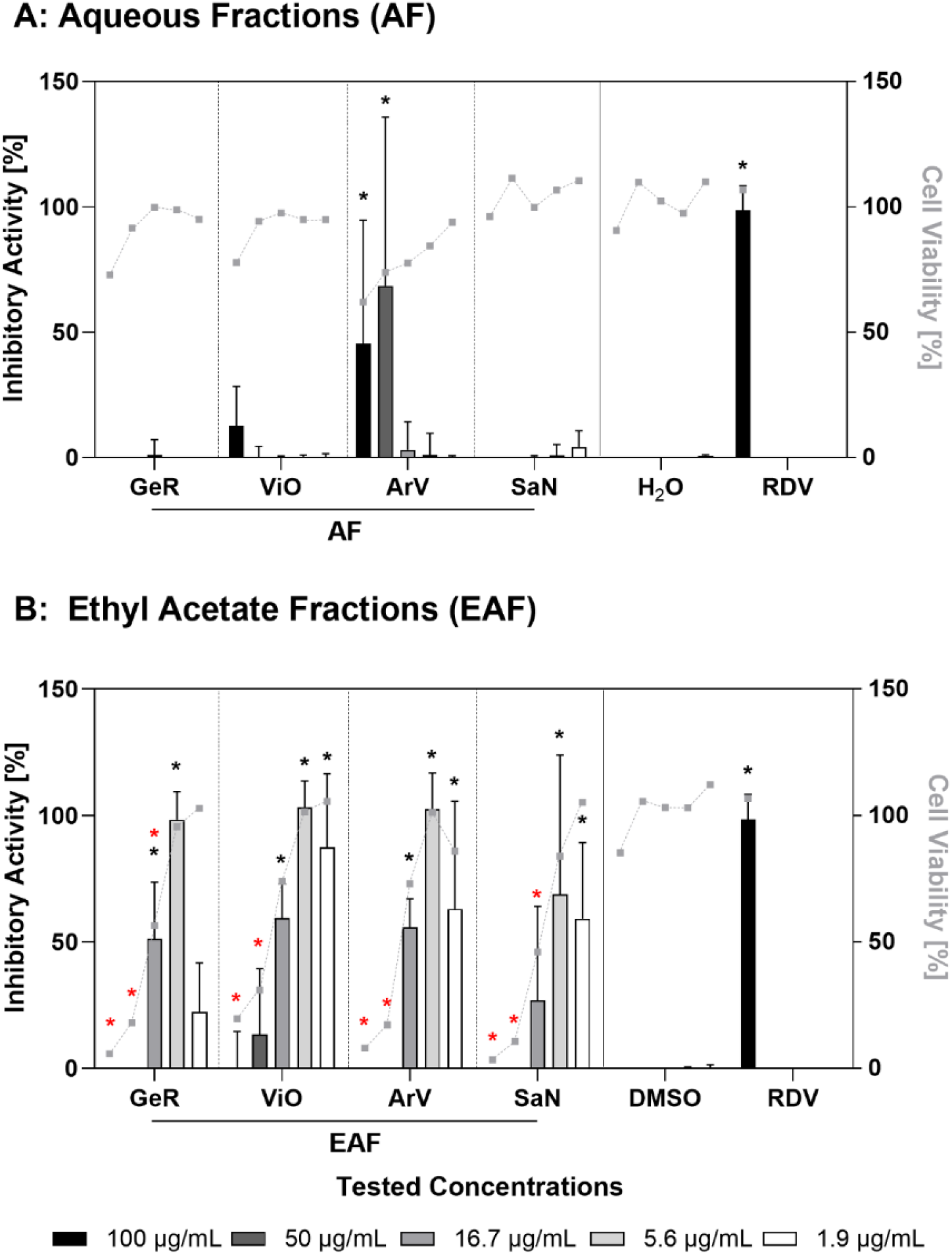
Inhibitory activity and cell viability of the ethyl acetate fractions. Inhibition of virus mediated cytopathic effect (bars) and absence of toxic effect (grey dots) of (A) aqueous fraction [AF] and (B) ethyl acetate fraction [EAF]. Abbreviation of tested plant species: GeR - Geranium robertianum, ViO - Viola odorata, ArV - Artemisia vulgaris, SaN - Sambucus nigra, H2O - solvent control aqueous fractions, DMSO - solvent control ethyl acetate fractions, RDV – Remdesivir as positive control. Means (n=3) of inhibitory activity and cell viability at individual concentrations were compared to their respective solvent control using two-way ANOVA with multiple comparisons corrected for using Dunnet’s method. Black asterisks (*).

### 3.4 HPLC-MS chemoprofiling

The EAF fractions of *Geranium robertianum* (aerial part) was further investigated by recording a metabolic profile using LC-PDA-HRESIMS (**Figure 3**). Main compounds found in the ethyl acetate fraction of *Geranium robertianum* are phenolic acids such as gallic acid (R_T_ = 1.08 min) and ellagic acid (R_T_ = 4.12 min) and flavonols such as quercetin (R_T_ = 8.25 min) and kaempferol (R_T_ = 10.19 min).

**Figure 3:**
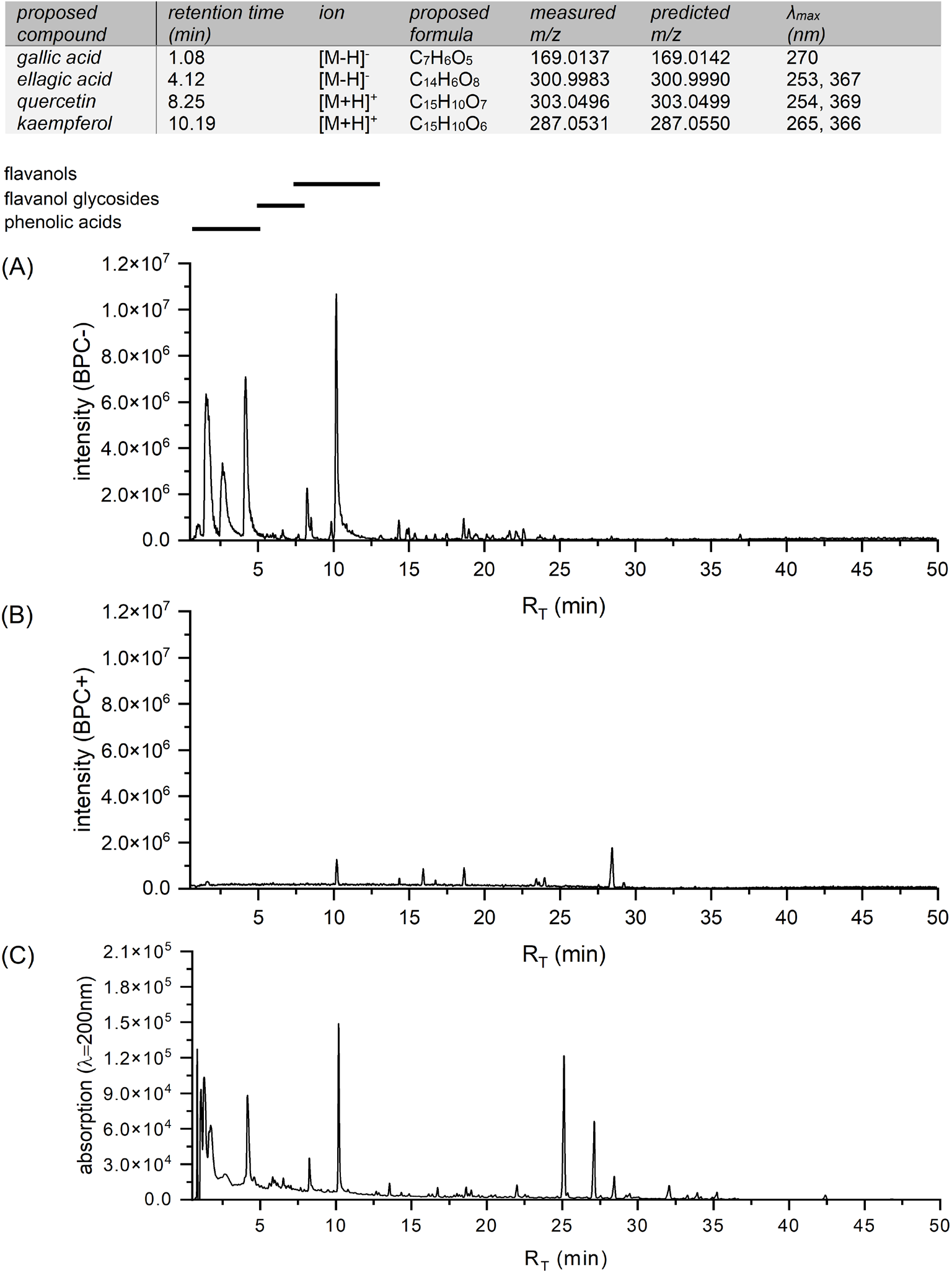
Metabolite profile of *Geranium robertianum* (aerial part) using LC-PDA-HRESIMS (A) base peak chromatogram (BPC) negative ionization mode (B) BPC positive ionization mode (C) chromatogram UV detection at λ=200 nm.

## 4 Discussion

All ten plant species selected from the *Receptarium* of Burkhard von Hallwyl and investigated in this study are native to the Swiss flora and have a centuries-long history of medicinal use in Europe, sometimes dating back to Antiquity as in the case of *Melissa officinalis* L., *Quercus robur* L., and *Sambucus nigra* L. (Dal Cero et al., 2014).

As previously suggested by Buenz et al. (2004), our results support the notion that historical written sources can be a promising tool for identifying potential candidates for pharmacological investigations. Notably, 4 of the 10 plants selected (40 percent), display an interesting *in vitro* anti-SARS-CoV-2 effect.

Especially organic solvent-based extracts and fractions from *Viola odorata* L. and *Geranium robertianum* L. exhibit a distinct inhibitory activity. At the same time these examples illustrate that our approach allows us to identify plants or plant parts that have so far been little researched in connection with the coronavirus. In both plants, we observed a marked protective effect with the ethyl acetate fractions at a concentration of 5.6 µg/mL. At the lowest tested concentration of 1.9 µg/mL, the ethyl acetate fraction of *V. odorata* (leaves)showed a promising antiviral activity of more than 85% without compromising cell viability. Violaceae are known to possess numerous cyclotides as for example the cycloviolacin VY1 isolated from the whole plant of *Viola yedoensis* Makino Bot. Mag. (accepted name *Viola philippica* var. *philippica*) with a reported activity against influenza A H1N1 virus (Liu et al., 2014; Mammari et al., 2021). Recently cycloviolacin peptides were investigated also *in silico* for their anti-SARS-CoV-2 activity (Bhandu et al., 2023). Interestingly, in traditional Persian medicine, common violet flowers are recommended in pulmonary diseases for the treatment of cough, pneumonia and pleurisy (Qasemzadeh et al., 2015). In a randomized double-blind controlled trial assessing the effect of *Viola odorata* syrup concomitant to conventional protocol, faster recovery and alleviation of respiratory symptoms in patients positively tested for SARS-CoV-2 were observed (Adel Mehraban et al., 2023). While in *Viola odorata* mainly the flowers seem to have been investigated, and not the leaves as in our study, *Geranium robertianum*, to our knowledge, does not seem to have been explored at all in context with SARS-CoV-2. The main compounds we found in the ethyl acetate fraction of *Geranium robertianum* are phenolic acids such as ellagic acid and flavonols, which is in line with other studies (Graça et al., 2016). Ellagic acid is a natural substance reported to have an antiviral effect on human rhinoviruses (Park et al., 2014), influenza A virus (Pavlova et al., 2018), Zika virus (Acquadro et al., 2020), Human Immunodefiency Virus-1 (Promsong et al., 2018), hepatitis B virus (Shin et al., 2005) and Ebola virus (Cui et al., 2018). However, it remains to be investigated which compounds of *G. robertianum* might prove responsible for the anti-SARS-Cov-2 activity observed in this study.

Only three of the plant species in our sample appear to have so far been investigated specifically for their *in vitro* anti-SARS-CoV-2 activity, namely *Salvia officinalis* L. (Le-Trilling et al., 2022; Farzam et al., 2024), *Artemisia vulgaris* L. (Kazachinskaia, 2022), *Sambucus nigra* L. (Eggers et al., 2022; Leka et al., 2022).

Besides the observed antiviral activity, each of the three most potent candidate species – *A. vulgaris, V. odorata* and *G. robertianum* – were also reported to possess anti-inflammatory and immunomodulatory activities, as suggested by various *in vitro* studies (Koochek et al., 2003; Afsar et al., 2013; Catarino et al., 2017; Liu et al., 2022). Since herbal preparations frequently exhibit polyvalent and pleiotropic effects on host defence systems, further research could also be directed at exploring a potential adaptogenic effect of the organic fractions derived from *V. odorata* L., *G. robertianum* L., and *A. vulgaris* L., as shown for instance for *Andrographis paniculata* (Burm.F) Wall.ex Nees and *Withania somnifera* (L.) Dunal (Panossian and Brendler, 2020).

## 5 Conclusion

This study exemplifies the potential of using premodern written sources as a starting point for the identification of possible candidate plants for pharmacological investigations. Using a systematic approach to select pertinent medicinal plant uses from the historical text, ten plant species were identified of which three showed a promising bioactivity profile in our study.

Although vaccination is so far the most important preventive measure, the emergence of new circulating variants poses a constant risk of undermining the effectiveness of available vaccines. Complementary preventive measures, ideally at the local entrance points of the pathogen, could help to further reduce the risk of infection. In this context, natural products acting as viral entrance inhibitors might constitute a promising avenue. Further investigations of our candidate plants, in particular with regard to the pharmacological properties contributing to the observed antiviral effect, could shed light on potentially existing prophylactic effects against SARS-CoV-2.

## 7 Author Contributions

NV: Conceptualization, Data curation, Methodology, Formal analysis, Investigation, Writing – Original Draft, Writing – Review & Editing, Visualization, Supervision,

JS: Data curation, Formal analysis, Investigation, Methodology, Visualization, Writing – Review & Editing

CU: Conceptualization, investigation, visualization, writing - original draft, writing-review and editing

EW: Funding acquisition, conceptualization, formal analysis, writing-review and editing

YG: Methodology, Investigation, Writing YR: Methodology, Investigation

OE: Conceptualization, Methodology, Investigation, Writing – Original Draft, Writing – Review & Editing, Visualization, Supervision

AL: Conceptualization, Methodology, Formal analysis, Investigation, Resources, Writing – Original Draft, Writing – Review & Editing, Visualization, Supervision, Project administration, Funding acquisition

## 9 Acknowledgments

We thank Barbara Frei Haller from Pro Thesauro Sanitatis (PTS), 5706 Boniswil (Switzerland), for access to the full version of the RBH database. The author accepted manuscript of this article appears on bioRxiv, https://doi.org/10.1101/2024.02.09.579674. Open access funding provided by ZHAW Zurich University of Applied Sciences.

