## Supplemental Tables TS1-TS3 for "Historical texts as a potential resource for plant-derived natural products against SARS-CoV-2 – the example of the *Receptarium* of Burkhard III von Hallwyl from 16th century Switzerland"

**Table S1.** Plant material

| No. | Bulk sample ID | Plant material name | Plant part | Supplier | Supplier lot no. |
|---|---|---|---|---|---|
| 1 | 20220003 | Violae odoratae Folia conc | folium | DIXA AG | 201667 |
| 2 | 20220004 | Querci Folia conc | folium | DIXA AG | 202156 |
| 3 | 20220005 | Sambuci Folia conc | folium | DIXA AG | 203084 |
| 4 | 20220007 | Salviae LMB Folia tot | folium | DIXA AG | 210630 |
| 5 | 20220008 | Alchemillae vulg. Herba conc | herba | DIXA AG | 211241 |
| 6 | 20220009 | Stellariae mediae Herba conc | herba | DIXA AG | 211824 |
| 7 | 20220010 | Geranium robertianum | herba | DIXA AG | 212663 |
| 8 | 20220011 | Veronicae Herba conc | herba | DIXA AG | 213169 |
| 9 | 20220012 | Violae odoratae Flos tot | flos | DIXA AG | 213691 |
| 10 | 20220014 | Melissae Folia tot elect | folium | DIXA AG | 214206 |
| 11 | 20220015 | Artemisiae Herba conc | herba | DIXA AG | 214542 |

**Table S2.** Solvents and consumables

| Article name | Supplier | Article no. | Supplier lot no. |
|---|---|---|---|
| Ethanol absolut | VWR | 20821.330 | 23B224007 |
| Ethyl acetate | VWR | 23882.321 | 22G294044 |
| Filter Paper | Macherey-Nagel | 729245.400 | 0.337 |

**Table S3.** Plants and extract preparations

| No. | Plant species | Plant part | Extraction medium | Drug-extract ratio | Extraction yield (%) [Dried extract weight / Dried plant material weight * 100] |
|---|---|---|---|---|---|
| 1 | *Viola odorata* L. | leaves | $H_2O$ | 5.3 | 19 |
| 2 | *Alchemilla vulgaris* L. | aerial part | $H_2O$ | 5.0 | 22 |
| 3 | *Geranium robertianum* L | aerial part | $H_2O$ | 4.3 | 24 |
| 4 | *Melissa officinalis* L | leaves | $H_2O$ | 3.4 | 29 |
| 5 | *Stellaria media* (L.) Vill. | aerial part | $H_2O$ | 3.9 | 25 |
| 6 | *Viola odorata* L. | flowers | $H_2O$ | 3.2 | 31 |
| 7 | *Artemisia vulgaris* L. | Aerial part | $H_2O$ | 4.7 | 21 |
| 8 | *Quercus robur* L. | leaves | $H_2O$ | 4.3 | 23 |
| 9 | *Salvia officinalis* L | leaves | $H_2O$ | 5.9 | 16 |
| 10 | *Sambucus nigra* L. | leaves | $H_2O$ | 3.9 | 26 |
| 11 | *Veronica officinalis* L. | aerial part | $H_2O$ | 4.0 | 25 |
| 12 | *Viola odorata* L. | leaves | EtOH 80% | 12.3 | 12 |
| 13 | *Alchemilla vulgaris* L. | aerial part | EtOH 80% | 8.5 | 12 |
| 14 | *Geranium robertianum* L | aerial part | EtOH 80% | 9.4 | 12 |
| 15 | *Melissa officinalis* L | leaves | EtOH 80% | 9.1 | 18 |
| 16 | *Stellaria media* (L.) Vill. | aerial part | EtOH 80% | 10.5 | 14 |
| 17 | *Viola odorata* L. | flowers | EtOH 80% | 5.5 | 28 |
| 18 | *Artemisia vulgaris* L. | aerial part | EtOH 80% | 10.9 | 14 |
| 19 | *Quercus robur* L. | leaves | EtOH 80% | 6.8 | 22 |
| 20 | *Salvia officinalis* L | leaves | EtOH 80% | 9.7 | 16 |
| 21 | *Sambucus nigra* L | leaves | EtOH 80% | 7.7 | 20 |
| 22 | *Veronica officinalis* L. | aerial part | EtOH 80% | 7.4 | 20 |
